# Spatiotemporal Transcriptomes of Pig Hearts Reveal Midkine-Mediated Vascularization in a Chronic Myocardial Infarcted Model

**DOI:** 10.1101/2023.06.10.544480

**Authors:** Swarnaseetha Adusumalli, Samantha Lim, Vincent Ren, Li Yen Chong, Clarissa Tan, Roy Tham, Ye Lei, Yibin Wang, Enrico Petretto, Karl Tryggvason, Lynn Yap

## Abstract

Ischemic heart disease is the most prevalent cause of death globally. Regenerative cardiology using stem cell-based therapy is a potential approach to replace infarcted myocardial (MI) heart tissue. We used cardiovascular progenitors (CVPs) derived from human pluripotent embryonic stem cells differentiated to cardiomyocyte progenitors on a laminin 521+221 matrix and transplanted them into acute and chronic MI pig hearts (AMI and CMI). We performed time-series spatial transcriptomics to characterize these human cells at AMI 1- and 2- and at CMI 1-, 4- and 12 weeks post-transplantation. Both models showed high transcriptional reproducibility in the replicates. Furthermore, the human grafts engrafted well, matured, and expressed metabolic, ribosomal, T-tubule, and channel-related genes in the human graft over time. Cell-cell communication analysis revealed Midkine (MDK) signaling as a key pathway that may lead to increased angiogenesis of collaterals in the human graft.

## INTRODUCTION

Cardiovascular diseases (CVDs) are a major health concern worldwide, with an increasing annual incidence of over 17% since 2010^1^. Within CVD, Ischemic heart disease (IHD) is the most prevalent cause of death. It develops due to the obstruction of blood flow to the myocardium, either by partial or complete blockage of a coronary artery leading to heart failure and reduced ejection fraction (HFrEF)^2^. The heart muscle becomes ischemic and partially necrotic due to the limited blood supply. The ischemic heart muscle does not regenerate which eventually leads to heart failure^2^. Regenerative cardiology using stem cell-derived cells is a promising strategy to replace damaged cardiomyocytes (CM) and improve heart function^3^. There is, however, limited understanding regarding the *in vivo* repair mechanisms of transplanted CMs cells in the development of successful approaches of regenerative cardiology.

The pathophysiological mechanisms driving IHD have been extensively studied in small and large animal models that mimic the complexities of CVD ^4^. Small animals such as mice, rats, and rabbits are frequently used in research because of their small size, inexpensiveness, ease of handling, and maintenance. However, these animal model system does not recapitulate the human heart due to its small heart size, fast heart rate, and anatomical differences in the coronary artery system^5^. In contrast, large animal models such as pigs, dogs, sheep, and monkeys resemble the human heart in terms of coronary collaterals, overall anatomy and physiology^6, 7^, as well as regeneration characteristics after a myocardial infarction (MI) injury^8^. Therefore, large animal models are widely used in studies of human genetic and cardiovascular diseases.

There are currently two types of MI animal models^9^, acute myocardial infarction (AMI) and chronic myocardial infarction (CMI) model. AMI is usually accomplished by coronary artery ligation or interventional embolization that leads to ischemic heart failure in the short term with active inflammation and ventricular remodeling, while CMI usually develops after the acute phase when scarring and fibrosis have developed in the ischemic region^9^. The CMI model simulates the natural pathogenesis of IHD and more closely resembles clinical manifestations in patients than the AMI model.

We have previously used both mouse and pig models in our studies to study the consequences of cellular regenerative cardiology^10, 11^. We have shown that cardiovascular progenitors (CVPs) derived from human pluripotent stem cells (hPSC) are able to reproducibly differentiate on a LN521+221 matrix and generate well-structured cardiac fibers in mouse and pig hearts^10, 11^. In addition, we have demonstrated the *in vivo* effects of CVPs on heart function improvement and reduction of ventricular arrhythmia^10^. In the present study, we further sought to investigate and characterize transplanted human CVPs in both AMI and CMI pig models. We performed time series spatial transcriptomics (ST) to evaluate the human cells in the 1- and 2-weeks AMI model, and 1-, 4- and 12-weeks in the CMI model. The ST analyses demonstrated successful transplantation of human CVPs both the AMI and CMI models. Subsequently, we utilized the CMI model for further analysis of long-term transplantation and graft maturation. In addition, we identified the human spot communications at the regions of transplantation which revealed MDK signaling as one of the key pathways that could may potentially lead to an increased vascularization in the human graft. In summary, our results provide a comprehensive spatial-temporal characterization of engrafted human cells in the CMI pig model that may reveal the potential mechanism of graft-induced angiogenesis.

## MATERIALS AND METHODS

### Pluripotent Stem Cells and CVP Differentiation

Pluripotent human embryonic stem cells (hESCs, H1) were purchased from WiCell Research Institute and studied with approval from the National University of Singapore Institutional Review Board (IRB 12-451). The protocol is based on our previous studies^10, 11^. Wild-type H1 cells were transduced with lentivirus particles carrying a luciferase-GFP-puromycin construct (Perkin Elmer, CLS960002) to generate an H1- luciferase cell line. The pluripotent H1-luciferase cells were regularly cultured on a 10 mg/ml LN521 matrix (Biolamina AB), in Nutristem medium (Biological Industries, 05-100- 1A). At differentiation day 0, the H1-luciferase cells were seeded at 6 million density in a 10 cm^2^ dish (ThermoFisher 150464). The dishes were pre-coated with LN521 (21.75 mg) and LN221 (65.25 mg) at a 1:3 ratio. At confluence (day 4), 10 mM of CHIR99021 (Tocris, 4423) were added into the differentiation medium (RMPI 1640 (ThermoFisher, 11875- 093) supplemented with B27 without insulin (ThermoFisher, A1895601). The next day, the medium was replaced with a fresh differentiation medium (without the addition of CHIR99021). On day 7, the medium was changed to a differentiation medium supplemented with 5 mM of IWP2 (Tocris, 3533) for 48 hours and changed to a new differentiation medium (without the addition of IWP2). Subsequently, the fresh medium was replaced every other day until the cells reached CVP (days 10 to 11) stage. The CVPs were cryopreserved with Cryostor CS-10 (Stem Cell Technologies, 07930) and stored in liquid nitrogen until the day of cell transplantation.

### Acute and Chronic Pig MI Model

All the animal experiments were performed with prior approval from Singhealth’s Institutional Animal Care and Use Committee (IACUC) (2018/SHS/1426). The sus scrofa pigs weighing 13–15 kg at approximately three months age were purchased from Singhealth Experimental Medicine Centre (Singapore).

The pigs were randomly allocated into three groups, CVP-transplanted, RPMI medium- transplanted control, and sham (healthy control). MI and CVP transplantation were performed on 2 pigs per time point for each MI model. For AMI, we investigated 1 and 2 weeks post-CVP transplantation, and for CMI, we investigated 1, 4, and 12 weeks post- CVP transplantation.

Detailed protocol for the AMI model (1 and 2-weeks) has been published previously^10^. Briefly, the pigs were immunosuppressed with daily oral administration of cyclosporine (Novartis, 15 mg/kg ADP835296) five days before surgery and intravenous administration of 12.5 mg/kg abatacept (Orencia, Bristol-Myers Squibb, ABT4318) on the day of surgery and every other week until the end of the experiment. On the day of MI surgery, we permanently ligated the first branch of the left anterior descending coronary (LAD) and left circumflex (LCX) arteries. Immediately after ligation, 200 million CVPs were injected intramyocardially into the ischemic region. The pigs were then allowed to recover and post-surgery analgesia and antibiotics (Ketoprofen, 5 mg/kg/day, Enrofloxacin 15 mg/kg) were administered for three days post-surgery. At 1- and 2-weeks post-transplantation, the pigs were euthanized, and D-luciferin (Perkin Elmer, 760505) was administrated into the left ventricles. The luciferase signal was observed by *in vivo* IVIS imaging. Areas with positive signals were sectioned for histological staining and visium spatial gene expression tissue processing (10x Genomics). Refer to Figure 1B for illustration.

**Figure 1.**
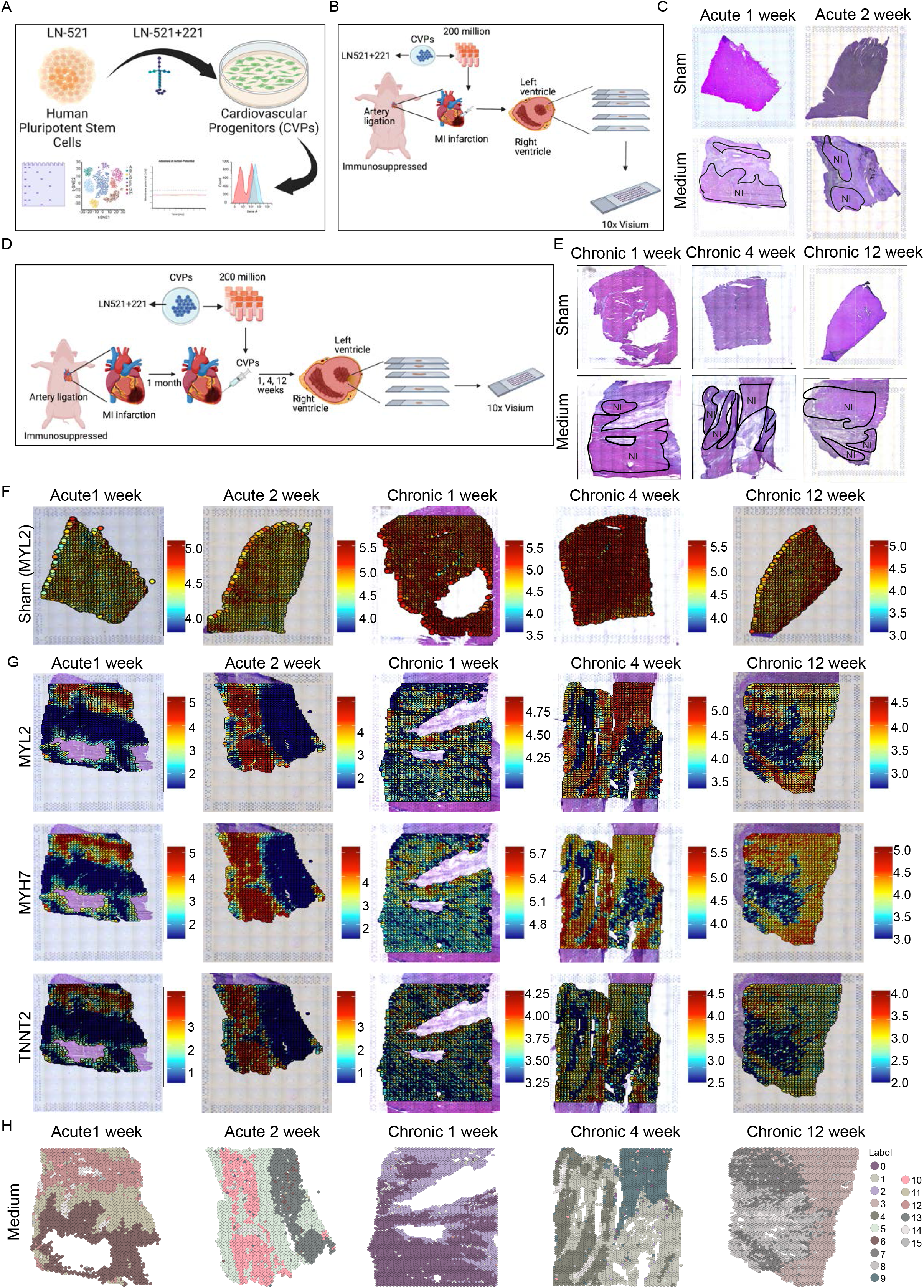
Spatial Transcriptomics of Acute and Chronic Myocardial Infarction Model in Pigs. **A)** Illustration of human pluripotent stem cells (hPSC) derived cardiovascular progenitors (CVPs) differentiated on laminin-221 (LN221) matrix. The CVPs were characterized using flow cytometry, electrophysiology measurements, single-cell RNA sequencing, and immunoblotting^10^. **(B)** Schematic of the CVP preparation, transplantation, and acute MI model creation in pigs. Left ventricles at 1 and 2 weeks were harvested and analyzed with 10x spatial transcriptomics. **(C)** H&E images of sham and medium control left ventricle in acute MI model. Non-infarcted (NI) region was demarcated with a black line while the rest of the tissue is the infarcted region. **(D)** Schematic of the CVP preparation, transplantation, and 1-month chronic MI model creation in pigs. Left ventricles at 1, 4, and 2 weeks were harvested and analyzed with 10x spatial transcriptomics. **(E)** H&E images of sham and medium control left ventricle in chronic MI model. Non-infarcted (NI) region was demarcated with a black line while the rest of the tissue is the infarcted region. H&E images with spots overlay in acute and chronic MI depict the normalized expression for **(F)** pig cardiac gene MYL2 in the sham tissue section and **(G)** human *MYL2, MYH7,* and *TNNT2* marker genes in medium tissue sections. **(H)** Spotplots with unbiased clustering based on global gene expression within individual spots in acute and chronic medium tissue sections.

For the CMI model (at 1, 4, and 12- weeks), the same coronary vessel ligation was performed on the pigs 4 weeks before the cell transplantation and either 200 million CVPs or RPMI medium were transplanted into the ischemic heart region. Similarly, the pigs underwent the same immunosuppression regimen 5 days before the cell transplantation, post-operation medications, and IVIS imaging. Refer to Figure 1D for illustration.

### Visium Spatial Expression Workflow

After the euthanization of the pigs, D-luciferin (15 mg/ml in 2ml volume) (Perkin Elmer, 122799-5) was administered into the excised whole heart via injection into both coronary arteries for 10 minutes. Subsequently, the left ventricle was isolated and cross-sectioned into 5 rings. All 5 rings were placed on the IVIS Spectrum imaging platform (Perkin Elmer) and 2D bioluminescent images were taken to identify areas with positive signals. These areas were snapped-frozen in liquid nitrogen and processed according to 10x genomics protocol (https://www.10xgenomics.com/support/spatial-gene-expression-fresh-frozen). Healthy and medium control tissue sections were similarly prepared for 10x visium workflow. After optimizing the permeabilization time with the pig heart tissue, we established concluded that 18 minutes was an optimal time to achieve the best images. RNA libraries were constructed and they underwent NovaSeq PE150 sequencing by Novogene AIT.

### Immunohistology Staining

Serial cryosections from the snap-frozen tissues were stained with anti-Ku80 (Cell Signaling, 2180S, RRID:AB_2218736, 1:300), anti-ACTN2 (Sigma, A2172, RRID:AB_476695 1:500) or anti-CD31 (Abcam, ab28364, RRID:AB_726362 1:100) at 4°C overnight. The next day, sections were washed thrice with 1 X PBS for 5 mins each. Alexa-conjugated secondary antibody (ThermoFisher, 1:1000) and DAPI (ThermoFisher, D1306, RRID:AB_2629482 1:5000) were added for 1 hr and washed thrice with 1 X PBS for 5 mins each. To suppress autofluorescence, the sections were incubated with Sudan Black (Sigma, 199664) for 20 mins, washed thrice with 1 X PBS, and then mounted with ProLong Gold antifade mountant (ThermoFisher, P36930). Slides were examined using a fluorescence microscope.

### Spatial Transcriptomics Data Processing

Time series spatial transcriptomes (ST) datasets were generated from pig heart left ventricles at 1- and 2- weeks for AMI and 1-, 4-, 12- weeks for CMI models. For each time point in both models, we have 4 tissue sections, sham, medium, and 2 replicates with human CVPs transplanted (i.e., a total of 20 tissues. 4 tissues x 2-time points in the AMI model and 4 tissue sections x 3-time points in the CMI model). Reads from the sham and medium (10 tissues) sections across each AMI and CMI week were mapped to the pig genome (Ensembl version 102) using a 10x space ranger (v1.2.1).

The remaining 10 tissue sections with human cells transplanted have reads from both humans and pigs. Therefore, to segregate the ST reads belonging to the transplanted human cells from the host pig heart reads, we prepared a combined genome reference from human and pig ensembl genomes (GRCh38 and Ssus11, v102) using *mkref* from spaceranger. The raw reads were aligned to the combined reference genome and the summarized UMI counts for each spot on the visium spatial transcriptomics array were generated using spaceranger *counts*. The mRNA count matrices were generated by adding intronic and exonic reads for each gene in each location. The paired histology H&E images were processed using spaceRanger to select locations covered by tissue by aligning to spot locations with fiducial border spots in the histology image. This enabled overlay of the quantified counts for genes in each spot. Only the spots under the tissue sections were retained for the downstream analyses.

The downstream processing was split into three separate analyses. First, the combined analysis of AMI and CMI model medium tissue sections. Second, analysis of replicates from 1-, and 2- weeks post CVPs transplanted tissue sections in the AMI model. Third, replicates from 1-, 4-, and 12- weeks post-transplanted human CVPs in the CMI model were processed. In each analysis, raw counts, images, spot-image coordinates, and scale factors were imported into the STutility R package (v1.1.1)^12^. The resulting gene-spot matrix generated was analyzed with the STutility and Seurat (v4.3) packages. As a quality control (QC), we removed genes that are expressed in less than 1 % of spots and less than 1 % of total reads. Spots expressing less than 5 % of the total number of genes were removed. Spots with more than 10 % of their total gene count coming from mitochondrial genes were also discarded. The resultant gene-spot information post-QC was summarized in Supplementary Table 1A. The post-QC gene-spot matrix counts were normalized across spots using regularised negative binomial regression (*SCTransform* function from Seurat package). Normalization and scaling across spots were performed by regression of the number of genes per spot.

Dimensionality reduction was performed using non-negative matrix factorization (*RunNMF* function in STUtility). The number of factors for the factor analysis was empirically set (16 factors for medium sections, 15 for AMI transplanted tissues, and 14 for CMI transplanted tissues) based on the spatial patterns in each analysis. For the medium sections, we performed the clustering analysis based on the NMF factors using the Shared Nearest Neighbor (SNN) algorithm using *FindNeighbors* and *FindClusters* functions from Seurat package at resolution 0.6. For AMI and CMI human CVPs transplanted tissues analyses, we identified the subgroups of spots specifically expressing human cells at the transplanted regions based on the NMF factors in two steps. Firstly, spots across tissues were clustered at resolutions 0.3 and 0.4 for AMI and CMI transplanted tissues, respectively. Identified clusters in each section were visualized in spatial context by overlaying spot over H&E images (with spot size scaling) using *ST.FeaturePlot* function from STutility. To identify markers within each cluster, we compared the cluster versus all others using the *FindAllMarkers* function (min. pct = 0.1, adjusted *P*-values <0.05 non-parametric Wilcoxon rank sum test). The number of human and pig genes expressed was computed per cluster. Secondly, in each cluster per spot level gene expression was extracted to confirm the clusters with engrafted human cells Supplementary Table 1B. The clusters in which each spot has at least 1% human genes expressed were labeled as human CVPs transplanted clusters. We further overlayed the spatial location of these clusters with H&E staining for further confirmation. After identifying the human CVPs transplanted clusters the expression of human and pig marker genes was plotted using the *FeatureOverlay* function in STUtility. We quantified and compared the expression levels of pig and human CVP markers in all the spots from the transplanted clusters. The expression levels across species and time points were plotted using vlnplot from Seurat and ggbarplots. The statistical differences across the pig and human CVP markers and across time points were computed using the *t.test* R function.

### Functional Analyses

We performed functional analysis for differentially expressed genes in medium and CVPs replicate sections using *enrichKEGG* and *gseKEGG* from ClusterProfiler R package (v3.18.1)^13^. The enriched pathways with adjusted *P*-value below 0.05 were plotted using *ggpubr* and *ggplot* (ggplot2.tidyverse.org). To account for reproducibility across the replicates, we computed Spearman’s ranked correlation of pig and human genes. The correlation across the replicates was visualized using the *pairs* R function.

### Cell-cell communication in transplanted spots

To investigate the signaling communication at each time point, independent spot clustering analyses were performed on each CMI time point individually. The low-quality reads and genes in each spot were filtered and spots with 10% of their total gene count coming from mitochondrial genes were also discarded. The replicates per timepoint were integrated using *FindIntegrationAnchors* and *IntegrateData*. The gene-spot counts from each timepoint were normalized by regularized negative binomial regression by using the function *SCTransform* with the default number of variable genes. 50 principal components were computed considering variable genes with the function *RunPCA*. Then the clustering analysis was performed using the Shared Nearest Neighbor (SNN) algorithm using *FindNeighbors* and *FindClusters* functions. The spatial spot clusters were plotted using *SpatialDimPlot*. UMAP was run using the function *RunUMAP* and clusters were plotted in UMAP space.

The ligand-receptor interactions and signaling in the human CVPs transplanted spot clusters at each CMI timepoint were predicted using CellChat^14^. The gene-spot counts, clusters, spot-image coordinates, and scale factors were fed to CellChat to identify communication signals in the form of ligand-receptor (LR) interactions *(P < 0.05)*. The number of interactions, their strengths, and the individual ligand-receptor pairs per timepoint were plotted using *netVisual_circle* and *netVisual_bubble* plot functions from CellChat. The shortlisted signaling circle plot was generated using *netVisual_aggregate* and the relative contributions of each LR pair were plotted using *netAnalysis_contribution*. The expression and spatial location of identified signaling genes were illustrated using bar charts, violin, and spot plots using ggbarplots, vlnplot from Seurat, and *FeatureOverlay* function from STUtility. Statistical significance was computed using the *t.test* R function.

### Statistical analysis

Comparisons between groups were performed using a t-test to find the significance of expression in the engrafted spots across species and across time points. Values were reported as Mean ± SEM. Replicate information is indicated in the figure legends and methods. A p-value < 0.05 was considered statistically significant. * p < 0.05.

### Data Availability

The ST data is made available on Gene Expression Omnibus. The non-sequencing data and materials are available from the corresponding author upon reasonable request.

## RESULTS

### Spatial Transcriptomics of Acute and Chronic Myocardial Infarction Model in Pigs

We have previously demonstrated high reproducibility of the differentiation protocol using the LN-521+221 matrix to differentiate hESC into well-characterized CVPs as confirmed with flow cytometry, electrophysiology measurements, scRNA sequencing, and immunoblotting analyses (**Figure 1A**). These CVPs improved heart function and reduced ventricular tachyarrhythmia in mouse and pig models^10, 15^. To further visualize and interrogate the transplanted human cells in animal models, we investigated the transplantation of the CVPs in AMI and CMI pig models by time-series spatial transcriptomics.

For the AMI model we captured two-time points post-transplantation (1- and 2 weeks, 2 replicates each), whereas in the CMI model, we captured three-time points post-transplantation (1-, 4- and 12 weeks, 2 replicates each) (**Figures 1B and D**). To generate both models, MI was induced by permanently ligating the coronary arteries either with immediate cell transplantation (AMI model) or transplanting cells after 1 month of ligation (CMI model) into the infarcted area. A healthy pig heart was used as a sham control and pigs transplanted with medium only was included as a negative control. H&E staining was performed on the sham and medium tissue sections to determine the area of non- infraction (NI) and infarction (**Figures 1C and E**). As expected, homogenous H&E staining was observed in the sham control indicating healthy non-infarcted tissue (NI), whereas, in the medium control tissue, fibrotic-like morphology was observed in various regions indicating MI within healthy NI tissue. This suggested that the coronary ligation was successful and that it resulted in heart infarction. These stained sections will aid in the validation of the downstream transcriptomic analysis.

The ST reads from healthy and medium control from both AMI and CMI were aligned to the pig reference genome and subjected to downstream computational analysis. To confirm MI models, we accessed the expression of key cardiac marker pig genes in sham and medium tissue sections across time. We observed the expression of the ventricular cardiac gene *MYL2* in all the spots in the sham healthy tissue (**Figure 1F**). On the other hand, we observed in the medium tissue specific spatial expression of cardiac markers *MYL2, MYH7,* and *TNNT2* in the NI region, but not in the infracted region as highlighted previously by H&E staining (**Figure 1G**). These results validated our ST pipeline showing that we can segregate the NI and infarcted regions with precision.

Furthermore, to identify subgroups of spots with similar gene expression profiles in NI and infarcted regions, we performed unsupervised clustering of the data. Using a graph-based approach, we embedded the data into low-dimensional space via non-negative matrix factorization and performed clustering on both AMI and CMI medium sections. To characterize the clusters and assess their spatial organization, spots were projected on the H&E-stained tissue images. We identified a total of 16 spot clusters and out of these 16, 8 clusters were specific to the NI regions (clusters 6 and 15 at 1-week AMI, cluster 11 at 2-weeks AMI, cluster 0 at 1-week CMI, clusters 5 and 10 at 4-weeks CMI, and clusters 3, 9 at 12-weeks CMI). The other 8 clusters were associated with the spots in the infarcted regions (**Figure 1H**).

To characterize molecular functions and pathways within the NI and infarcted regions, differential gene expression analysis and gene set enrichment analysis were performed. We observed that the NI regions were highly enriched for genes associated with cardiac functions including cardiac muscle contraction, diabetic cardiomyopathy, and others in both AMI and CMI models (adjusted *P* < 0.05) (**Supplementary Figure 1A**). As expected in NI regions of the CMI model, we observed upregulation of metabolic pathways such as fructose, mannose, galactose, 2-oxocarboxylic acid, pentose phosphate, and glycolysis which are important pathways for a functioning heart muscle^16^ (Supplementary Figure 1A). In contrast, genes upregulated in infarcted regions were enriched for pathways such as protein digestion, endocytosis, ECM-receptor interaction, focal adhesion, and others which are associated with matrix remodeling are typical for an infarcted tissue^17^ (**Supplementary Figure 1B**).

### Identification of Human Cells Post-transplantation

After ligating the coronary arteries, CVPs were transplanted into the infarcted heart in duplicates of either AMI or CMI models. At various time points, left ventricles were harvested and sectioned for H&E and immunofluorescence staining (**Figures 2A and B**). In agreement with our previous study where we showed a significant increase in human graft size across time, we observed a larger graft at 12 weeks as compared to 1 week^10^.

**Figure 2.**
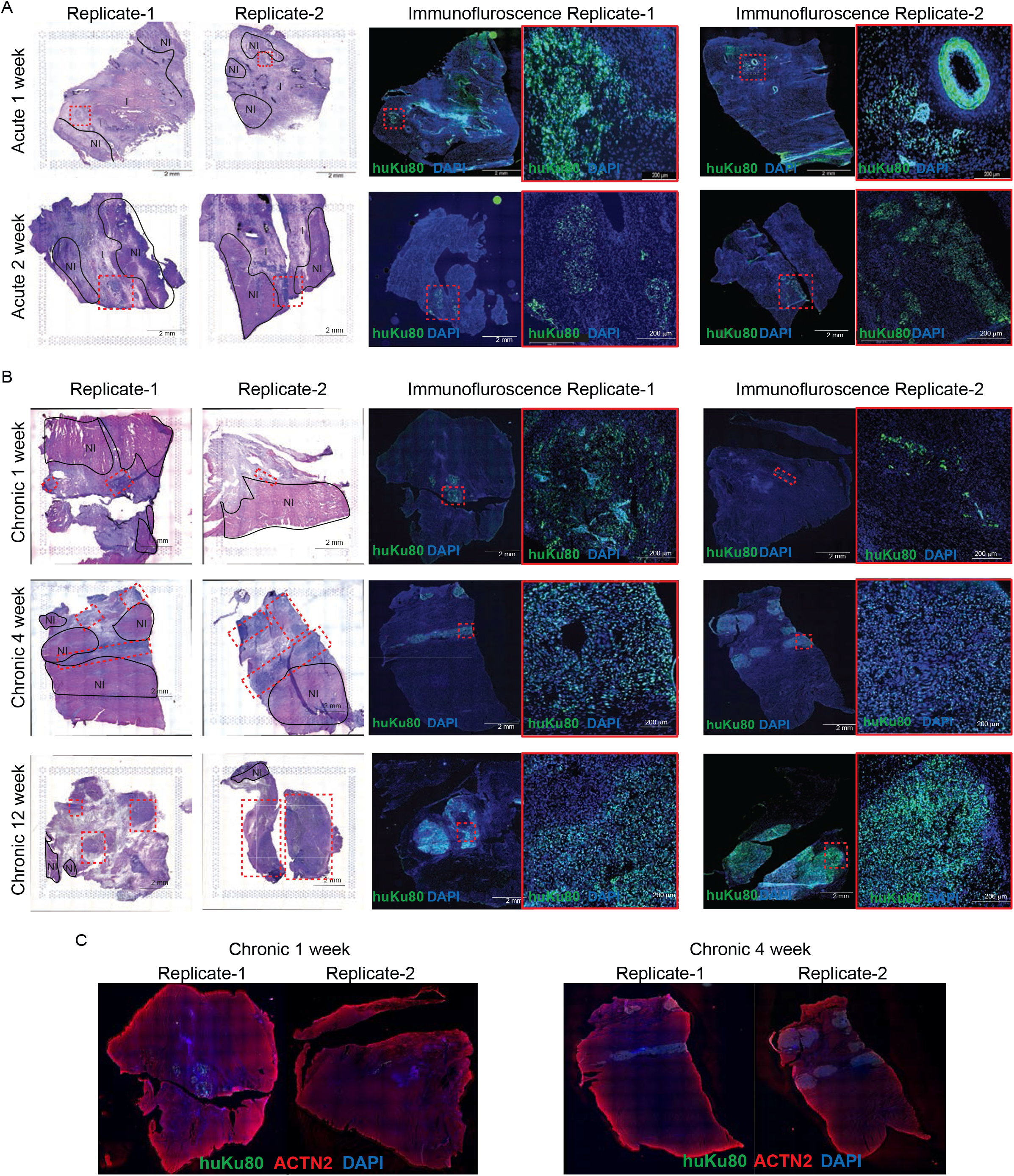
Identification of Human Cells Post-transplantation. Immunohistology staining in two biological replicates **(A)** acute MI at 1- and 2- weeks and **(B)** chronic MI at 1-,4- and 12 weeks post-transplantation. (Left) H&E images of both replicates and the engrafted human cells are highlighted regions in the red box. The non- infarcted regions in the tissues are demarcated with a black line (NI) and the rest of the tissues are infarcted (I) regions. (Right) Immunofluorescence images of engrafted human cells identified using human-specific Ku80 (huKu80) antibody (green). The nucleus is stained with DAPI (blue). Insert shows the higher magnification of engrafted human cells from the dotted red box.

Human-specific anti-Ku80 (green) was used to identify and visualize the human graft in the infarcted region. Human grafts were highlighted in dotted red boxes and a magnified view of the engrafted region was shown in the right panel. Following that, we stained the tissues with cardiac-specific anti-alpha actinin 2 (ACTN2) (red) to confirm the presence of human cardiac cells (**Figure 2C**). These data provided the ground truth for the exact location of the transplanted human cells and where our ST pipeline will reference it to.

### Characterization of Engrafted Human Cells in AMI Pig Hearts

The 1- and 2-week AMI left ventricles were processed in spatial transcriptomic workflow and the reads from the post-transplanted tissue sections (replicate-1 and replicate-2) were aligned to the pig and human combined genome references to segregate the reads belonging to the transplanted human cells from the pig. After mapping and quality control, we obtained individual gene-spot matrices summarized over four sections at each week (**Supplementary Table 1A**).

To confirm the successful transplantation of human cells, we assessed the expression of human cardiac markers *ACTC1, ACTN2, MYL2, NKX2-5, TNNT2,* and *TPM1* in the tissue sections of replicate 1 (**Figures 3A-F**). Supplementary Figures 2A to F show the results from the same genes but for tissue sections of replicate 2. We showed that these human markers were indeed specifically expressed in the same spatial region where human cells are as highlighted in Figure 2. On the contrary, when we assessed the same genes but from pig origin, results showed that in the same human spatial region, we measured a low expression of these pigs’ genes. This strongly suggests successful transplantation and engraftment of CVPs into an infarcted pig heart.

**Figure 3.**
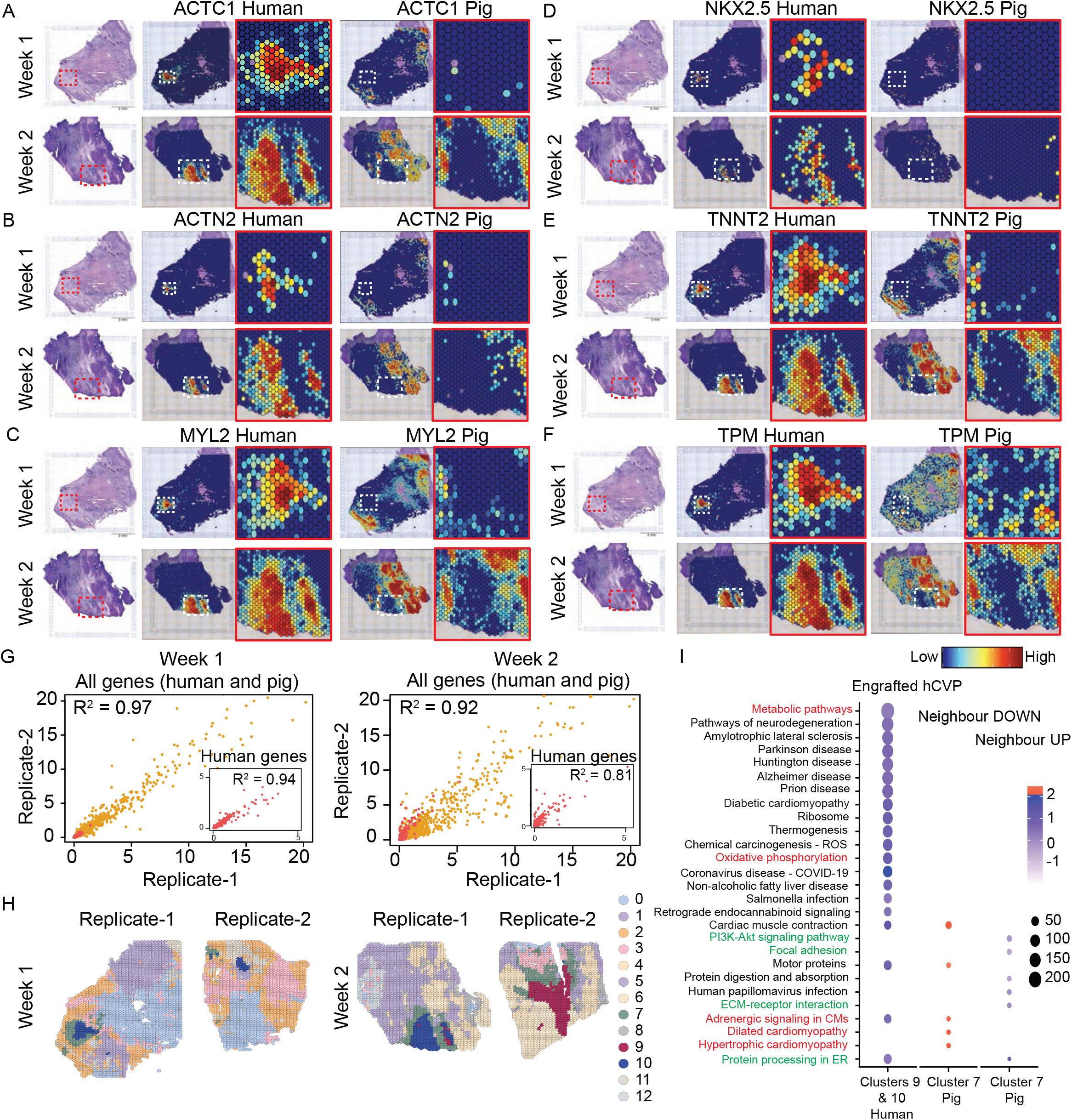
Characterization of Engrafted Human Cells in AMI Pig Hearts Using Spatial Transcriptomics. **A-F)** H&E and spot overlays depict the normalized expression for human and pig *ACTC1, ACTN2, MYL2, NKX2-5*, *TNNT2,* and *TPM1* marker genes at 1- and 2-weeks post-transplantation in replicate-1. (Left) H&E images with engrafted human cells are highlighted in red. Insert shows the higher magnification of engrafted human cells from the dotted white box. **(G)** Correlation plot of all genes including pig and human genes from two replicates (R^2^ = 0.97 and 0.94 at 1- and 2-weeks). Insert shows the correlation of human genes only in the engrafted regions in both replicates (R^2^ = 0.94 and 0.81). **(H)** H&E and spot overlays with unbiased clustering of spots based on global gene expression within individual spots in 1- and 2- weeks post-transplantation. Human cells are identified in clusters 9 and 10 while immediate neighbouring pig spots are in cluster 7. **(I)** Functional pathways in human gene clusters 9 and 10. The down and up-regulated functional pathways from the immediate neighbouring pig cluster 7 were identified. Normalized enrichment scores (NES) denote the upregulation and downregulation enrichment strength (adjusted *P*-values < 0.05) and the size of the spots indicates the number of enriched genes. Highlighted in red are the cardiac pathways that were upregulated in the human cells cluster and downregulated in neighbouring pig cluster. Pathways in green color font were enriched in neighbouring pig clusters.

Subsequently, we quantified the expression levels of the pig and human markers in the transplanted regions each week (**Supplementary Figures 2G and H**). We observed significantly higher expression of human markers *ACTC1, MYL2, TPM1,* and *TNNT2* compared to those from pigs at 1- and 2- weeks post-transplantation (*P* < 0.05), whereas human *NKX2-5* was only significantly expressed in 2-week post transplanted tissue sections (**Supplementary Figures 2G and H**). The normalized expression levels of these marker genes in a violin plot across time points were shown in **Supplementary Figure 2I**. We then sought to assess the transcriptional similarity across both the CVPs transplanted replicates and performed correlation analysis across human and pig genes from the whole tissue sections as well as for human genes only in the transplanted regions at 1- (**Figure 3G, left)** and 2- weeks (**Figure 3G, right)**. The correlation analysis revealed high reproducibility between replicates at both time points (Spearman’s correlation, R^2^ levels of 0.97 for both human and pig genes and 0.94 for human genes only at 1- week, and 0.92 and 0.81 at 2- weeks post-transplantation respectively).

To further characterize spots with transplanted human cells, we identified the subgroup of spots specific to human cells in both replicates across time points in two steps. First, we performed unsupervised clustering to group the spots with similar gene expression profiles and identified the human clusters by a data-driven approach. This clustering analysis revealed a total of 13 spot clusters (**Figure 3H**). Second, we performed spot-level analysis to identify the clusters expressing at least 1% of human genes in each spot (**Supplementary Table 1B)**. The spot clustering and individual spot-level analysis revealed two clusters (clusters 9 and 10) accounting for most human genes and importantly our unbiased single-spot level analysis reliability predicted human clusters that most similarly represented the ground truth in Figure 2. Cluster 10 expressed human genes in both replicates from 1- week and replicate-1 from 2- weeks. Whereas cluster 9 accounts for the human genes in replicate-2 at 2- weeks post-transplantation (**Figure 3H**).

The differential gene expression analysis from the human clusters 9 and 10 in comparison with other clusters identified 1414 human genes (in 89 % of spots, n = 425) in cluster 9, and 1722 (in 97% of spots, n=321) human genes in cluster 10 respectively (**Supplementary Table 1C**). Of these, 1308 (93 %) genes were common at both time points, which includes cardiac markers such as myosin-chain genes, troponins, and myocardin (**Supplementary Table 1D**). We also observed about 15 % (n = 201) of human genes expressed to be mitochondrial (n = 119) and ribosomal (n = 82) genes as cells undergo mitochondrial biogenesis and activation of oxidative phosphorylation as key regulatory events in cell differentiation to beating CMs^18^. The remaining human genes include various genes such as collagens, laminins, heterogeneous nuclear ribonucleoproteins, and others.

Strikingly, we identified the human transplanted clusters across 1- and 2- weeks that have spatially shared their border with pig cluster 7. To further investigate the biological differences between these two neighbour clusters we performed a gene set enrichment analysis. Interestingly, the functional analysis revealed significant upregulation of cardiac pathways in the human CVPs transplanted clusters and downregulation of the same pathways in neighbouring pig clusters (adjusted *P* < 0.05) (**Figure 3I**). These pathways include metabolic pathways, oxidative phosphorylation, adrenergic signaling in cardiomyocytes, dilated cardiomyopathy, and hypertrophic cardiomyopathy which are pathways related to cardiac functions^18^ (highlighted in red, **Figure 3I**). Furthermore, the neighbouring pig cluster 7 showed upregulation of PI3K signaling, focal adhesion, ECM-receptor interaction, and protein processing which are related to ECM remodeling^19^ (highlighted in green, **Figure 3I**).

Taken together, using the AMI model we have shown successful transplantation of human cells in the infarcted host pig hearts. We showed significant expression of key human cardiac marker genes, such as *ACTC1, MYL2, TPM1,* and *TNNT2* at 1- and 2- weeks post-transplantation. We also demonstrated the high reproducibility of transplantation across biological replicates. Furthermore, we observed the up and down- regulation of the same cardiac pathways in the human CVPs transplanted cluster and the neighbouring pig cluster.

### Characterization of Engrafted Human Cells in CMI Pig Hearts

Similarly, to the AMI model, we utilized ST analysis in the CMI model by assessing the expression of key cardiac marker genes at 1-, 4- and 12- weeks post-transplantation. We observed specific expression of human marker genes *ACTC1, ACTN2, MYL2, NKX2-5, TNNT2,* and *TPM1* in the cells transplanted regions in replicate 1 (**Figures 4A to F).** These regions were previously shown to contain human cells by immunofluorescence staining in Figure 2. For replicate 2, the spot plots of these human markers are shown in Supplementary Figures 3A to F. We then quantified expression levels of these pig and human markers at the engrafted regions each week. At 1-week post-transplantation, there was a significantly higher expression of human cardiac markers *ACTC1, MYL2,* and *TPM1* compared to those from pigs in both replicates (*P* < 0.05). In contrast, at 4- and 12- weeks, all the human markers are significantly expressed in comparison with their pig orthologs (**Supplementary Figure 3G**). The normalized expression levels of these marker genes across time points were shown in **Supplementary Figure 3H**. Consistent with the AMI model, the correlation analysis in the CMI model suggested high transcriptional reproducibility across replicates at all time points (Spearman’s correlation, R^2^ levels of 0.96 and 0.8 for both human and pig genes and human genes at 1- week; 0.97 and 0.99 at 4-weeks, and 0.88 and 0.98 at 12-weeks post-transplantation respectively **Supplementary Figure 3I**).

**Figure 4.**
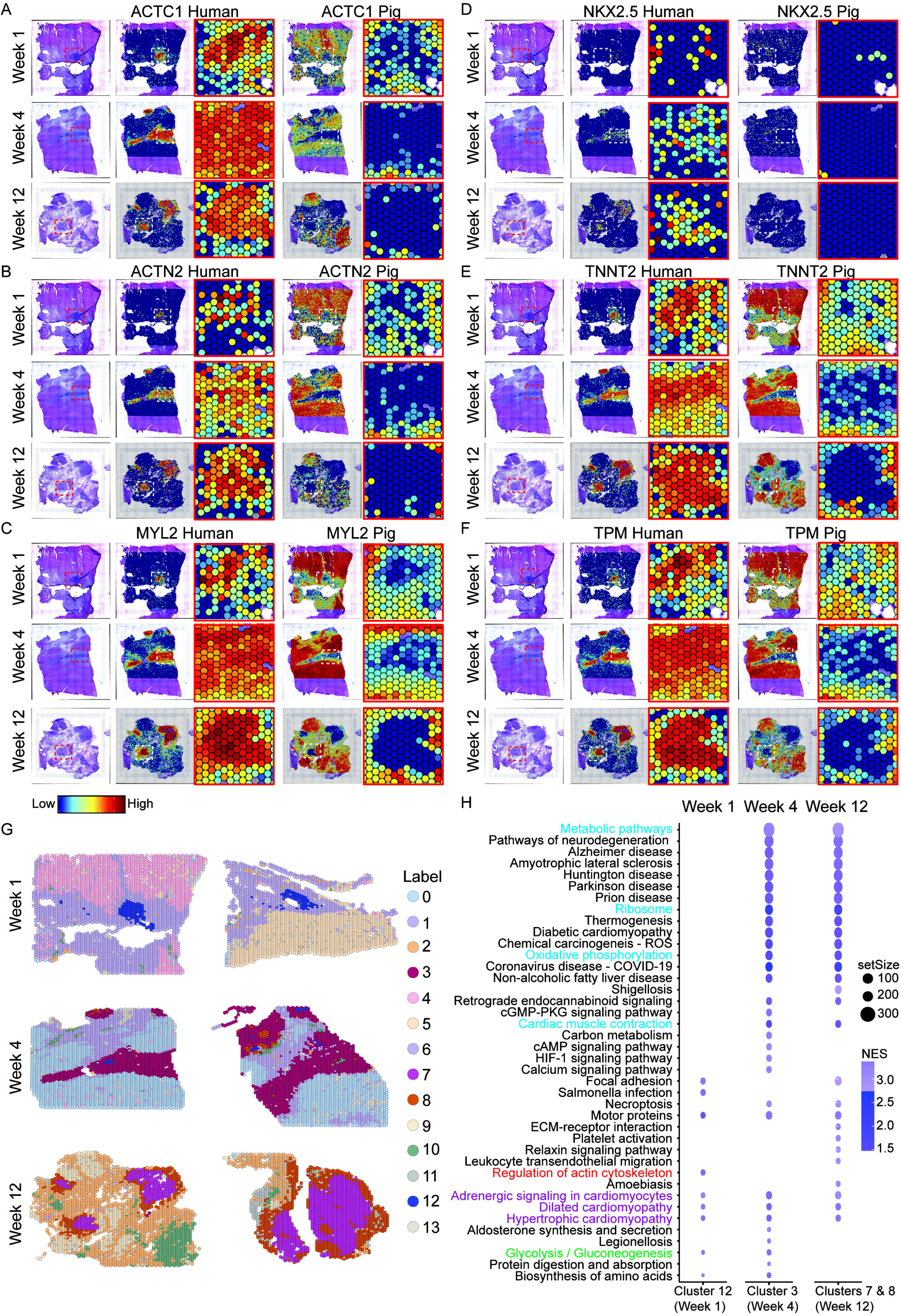
Characterization of Engrafted Human Cells in CMI Pig Hearts Using Spatial Transcriptomics. **A-F)** H&E and spot overlays depict the normalized expression for human and pig *ACTC1, ACTN2, MYL2, NKX2-5*, *TNNT2,* and *TPM1* marker genes at 1-, 4- and 12-weeks post- transplantation in replicate-1. (Left) H&E images with engrafted human cells are highlighted in red. Insert shows the higher magnification of engrafted human cells from the dotted white box. **(G)** H&E and spot overlays with unbiased clustering of spots based on global gene expression within individual spots in 1-, 4- and 12- weeks post-engrafted tissues. Human cells are identified in cluster 12 at 1- week, cluster 3 at 4- week, and clusters 7 and 8 at 12- week. **(H)** Functional pathways in human gene clusters at 1-, 4- and 12- weeks. Normalized enrichment scores (NES) denote the upregulation and downregulation enrichment strength (adjusted *P*-values < 0.05) and the size of the spots indicates the number of enriched genes. Pathways upregulated at 1- weeks were highlighted in red. Those enriched at and genes enriched at 1- and 4- weeks were in green. Pathways upregulated at 4- and 12- weeks were highlighted in blue, and cardiac common pathways were in purple.

Next, we identified the subgroup of spots specifically expressing human cells at the engrafted regions in two steps (**Figure 4G**). Firstly, we identified 13 spatiotemporal spot clusters and projected the clustered spots on the H&E-stained tissue sections. Secondly, we performed spot-level analysis to determine the human clusters with at least 1 % of human genes in each spot per cluster (**Supplementary Table 1B**). Clusters 12 and 3 spatially marked the regions of human cells at 1- and 4- weeks respectively while clusters 7 and 8 marked the human cells at 12- weeks (**Figure 4G**). The differential gene expression analysis identified 667 human genes in cluster 12 compared to all other clusters at 1- week post-transplantation (62 % of spots, n = 134). There were 2329 human genes in cluster 3 (74 % of spots, n = 1535) at 4- weeks and 3544 human genes in clusters 7 and 8 (100 % of cluster 7 spots, n = 1302 and 68 % of cluster 8 spots, n = 688) at 12- weeks (**Supplementary Table 1E**).

Furthermore, we compared the human DE genes from three-time points and identified 512 common genes that include troponins, myosin-chain, and others (**Supplementary Table 1F**). Similar to the AMI model, we observed about 18 % of common human DE genes expressing mitochondrial (n = 41) and ribosomal genes (n = 51). There were 69, 301, and 668 DE genes that were uniquely identified at time points 1-, 4- and 12- weeks. The functional enrichment analysis of human DE genes across time points revealed three common upregulated cardiac-related pathways: namely, adrenergic signaling, dilated cardiomyopathy, and hypertrophic cardiomyopathy (highlighted in purple, **Figure 4H**). We observed genes enriched for regulation of actin cytoskeleton at 1-weeks (highlighted in red) and genes enriched for glycolysis at 1- and 4- weeks (highlighted in green). Notably, the genes upregulated at 4- and 12- weeks were enriched for metabolic pathways, ribosomes, oxidative phosphorylation, and cardiac muscle contraction (highlighted in blue).

### Metabolic Maturation of Human Grafts

Previous studies have shown an increase in metabolic, mitochondrial biogenesis, ribosomal activity, and myofibril isoform switches such as MYH6 and MYH7 as some of the hallmarks of cardiomyocyte maturations^16, 20–22^. To understand the maturation status of the cells, we investigated these maturation hallmarks in our human graft across time from the CMI model.

We observed a significant increase (p < 0.05) in the expression of ribosomal and mitochondrial genes as well as the fuel consumption switch from immature glucose metabolism using glycolysis to a more mature fatty acid metabolism using oxidative phosphorylation across the weeks^16^ (**Supplementary Figures 4A-B and Figure 4H**). Furthermore, the significant increase in expression of the calcium and potassium channel genes (*CACNA1C, CACNAB2, RYR2, KCNH2, KCNJ8,* and *KCNMB4*) as well as gap junction connexin-43 (*GJA1*) that contributed to the formation of mature T-tubule structure and cell-cell communications^16^ indicating the maturing status of the human graft. We represented *GJA1* in a spot plot to display the spatial expression of this human gene in the engrafted region together with the absence of pig *GJA1* in the same location (**Supplementary Figure 4D**).

Other cardiac maturation markers such as *CD36*^20^ and *ESRRA*^23^ are also significantly increased over time (**Supplementary Figure 4C**). Interestingly, we observed an increase in the mature *MYH7* and a concomitant decrease in the immature *MYH6* expression which signify the MYH6/MYH7 structural protein switch of a maturing cardiac phenotype^16^ (**Supplementary Figure 4E**).

Taken together, these results suggested that the initial transplanted CVPs developed and differentiated into mature CM over time in the CMI model. The highly expressed metabolic, ribosomal, mitochondrial, and mature sarcomere proteins and the t-tubules structures are suggestive of a mature and functional human cardiac graft. Therefore, we are interested in elucidating the potential cell-to-cell communications in human grafts for further understanding of cardiac regeneration.

### Cell-cell Communications in Human-Engrafted Spots

While AMI is a life-threatening emergency, which benefits from early diagnosis and treatment, the prevalence of AMI is estimated between 1- 2 % in the general population^10^. In contrast, the global prevalence of chronic MI is 3.8 % in < 60-year-old patients and 9.5% in > 60-year-old patients. Due to the accelerated rate of MI in the aging population, our attention to CMI seems critical^24^. Having characterized both AMI and CMI models, we have chosen to focus on graft development in the CMI model because this model is more clinically relevant.

Studies have emphasized the application of cell-cell communication using ST data, as cell communications are spatially constrained^25, 26^. Understanding the cell-to-cell communication in the ST time course data will be useful to gain insights into cardiac regeneration. Therefore, we explored cell-to-cell communication by identifying and analyzing the conserved and time-specific signaling from the transplanted human cells in the CMI model. To study the signaling at each time point, spot clustering analyses were performed on each time point individually. We identified 11 spot clusters each at 1- and 4- weeks and 10 clusters at 12- weeks which were represented in the spatial context (**Figures 5A-C**) and Uniform Manifold Approximation and Projection (UMAP) (**Supplementary Figures 5A-C**). The differential gene expression analysis revealed human spot clusters (C7 at 1- week; C1, C5, C6 at 4- weeks; and C2, C5, C7, and C9 at 12- weeks) corresponding to the human cells transplanted regions as identified through H&E staining in Figure 2 (human cluster numbers are highlighted in red).

**Figure 5.**
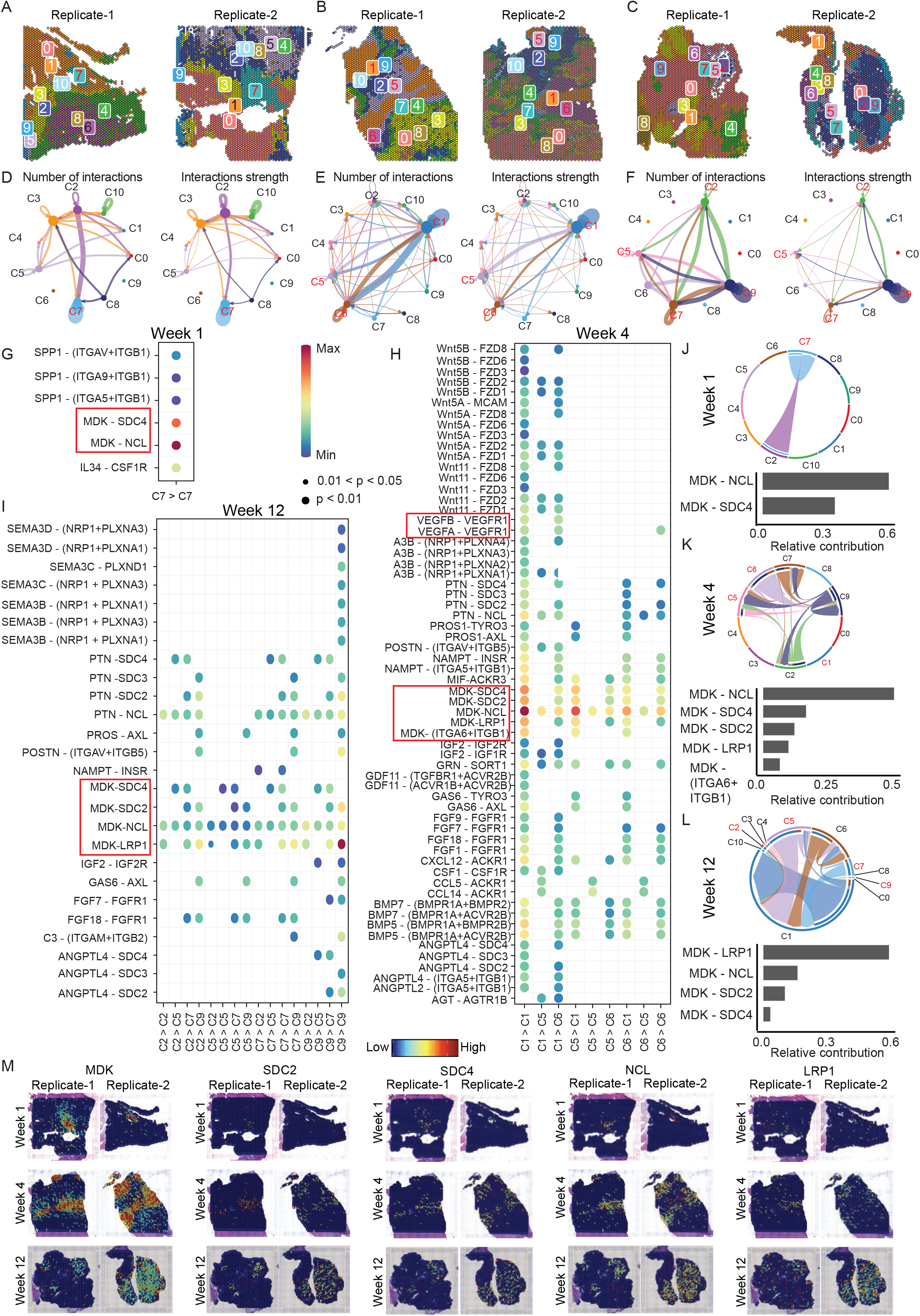
Cell-cell Communications in Human-Engrafted Spots. Spotplots showing the unsupervised clustering of spots based on global gene expression within individual spots for both replicates at **(A)** 1- week **(B)** 4- weeks and **(C)** 12- weeks post-transplantation. **(D-F)** Circle plot showing the number and strength of ligand-receptor interactions at each time point. Nodes and edges represent the ligand-receptor pairs. Node sizes and the edge line thicknesses are proportional to the number and the strength of the ligand-receptor signal. The arrows indicate incoming or outgoing signalling at each cluster. The round loops at clusters represent the interactions within the same cluster. At week 1, cluster 7 represents the human cells, at week 4, clusters 1, 2, 5, and 6 represent the human cells and at week 12, clusters 5, 7, and 9 represent human cells. **(G-I)** The bubble heatmap plot shows the ligand–receptor pairs contributing to signalling from one human cluster to other human clusters. The size of the dot indicates the p-value significance, and the color indicates the importance of the ligand-receptor pair in the corresponding network. **(J-L)** Circle plots showing MK signalling pathways network at each time point. The arrowheads in the plot indicate incoming or outgoing signalling at each cluster. The corresponding bar charts show the relative contributions of each ligand-receptor pair in MK signalling. **(M)** H&E and spot overlays depict the normalized expression for human MK signalling ligand and receptor genes *MDK, SDC2, SDC4, NCL,* and *LRP1* at 1-, 4- and 12-weeks post-transplantation.

The cell-to-cell communication network analysis was performed to identify statistically significant ligand-receptors interaction pairs for each time point (**Figures 5D-F, Supplementary Figure 5D-F**). We observed many strong interactions between and within the human clusters throughout the weeks. We further explored the subset of ligand receptors that were specific to the human-enriched spot clusters (**Figures 5G-I**). We observed SPP1- integrins (ITG) and interleukin-34 (IL34) - colony-stimulating factor 1 receptor (CSF1R) signaling were predominant at 1- week, suggesting cardiac fibroblast proliferation and tissue homeostasis^27–29^ (**Figure 5G**). Whereas at 4- weeks, signaling such as WNT^30^, VEGF^31^, FGF^32^, IGF^33^, and BMP^34^ were significantly enriched suggesting cardiac hypertrophy, development, and blood vessel formation (**Figure 5H**). At 12- weeks we observed specific signaling of semaphoring (SEMA^35^), complement component 3 (C3^36^) (**Figure 5I**) which suggested regulation in cardiac development and remodeling. It was also known that the gene periostin (*POSTN*^37, 38^) contributes to postnatal CM maturation, and innervation which could promote cardiac repair. Moreover, this gene is expressed at both weeks 4 and 12 suggesting cardiac regeneration (**Supplementary Figure 5G**).

Interestingly, we observed a common ligand Midkine (MDK) that was expressed throughout all the weeks with different cell membrane receptors (such as syndecans (SDC^39^), LDL receptor-related protein 1 (LRP1^40, 41^), nucleolin (NCL^42^) and integrins (ITG^27^)) (**Supplementary Figure 5H**). It was shown that the expression of MDK is beneficial to cardiac regeneration by improving CM survival after MI^43^. Moreover, its receptors like SDC4, NCL, and ITGB1 have protective effects on the heart from myocardial injury and myocardial proliferation. The relative contribution from MDK ligand-receptor interactions was plotted (**Figures 5J to L, Supplementary Figures 5H and I**). The MDK-NCL interaction contributed to the highest signaling at weeks 1 and 4 while MDK-LRP1 signaling is the highest at week 12. The spatial-temporal expression of these human genes was displayed in the spot plots and could visualize the expression of these genes at the human clusters (**Figure 5M**).

In summary, using cell-to-cell communication analysis, we have identified specific and common ligand-receptor interactions throughout the development of human grafts. Importantly, these interactions are involved in the normal cardiac development pathways.

### Expression of Vascularization-related genes

MDK was shown to be associated with cardiac pathology and angiogenesis^44^. It has been demonstrated that MDK is both necessary and sufficient to promote endothelial growth factor A cell proliferation in growing collaterals^45^. As angiogenesis is a well-studied process for organ development and regeneration especially in the heart where its main function is to pump blood throughout the body, important angiogenic factors such as vascular endothelial growth factor A (VEGFA) and vascular endothelial growth factor B (VEGFB). Therefore, we hypothesized that MDK expression might mediate angiogenic factors such as VEGFs in the MI heart tissues in the CMI model. To gain further insights, we have chosen to examine VEGFs further. Spot plots of human *VEGFA* and *VEGFB* showed a significant increase in expression from weeks 1 to 4 and no significant increase from weeks 4 to 12. (**Figure 6A and Supplementary Figure 5J**). This indicates that the stimulation of blood vessel formation peaks at week 4 and is maintained from week 4 to 12. Other than the expression of human *VEGFA* and *VEGFB*, angiogenic-related genes such as pig-origin von Willebrand factor (*VWF*) and *CD31* were also detected in the human transplanted region (**Figures 6B and C**). These data suggested that the human graft could be secreting potent VEGFA and VEGFB to stimulate the collateral formation from the host arteries into the engrafted region.

**Figure 6.**
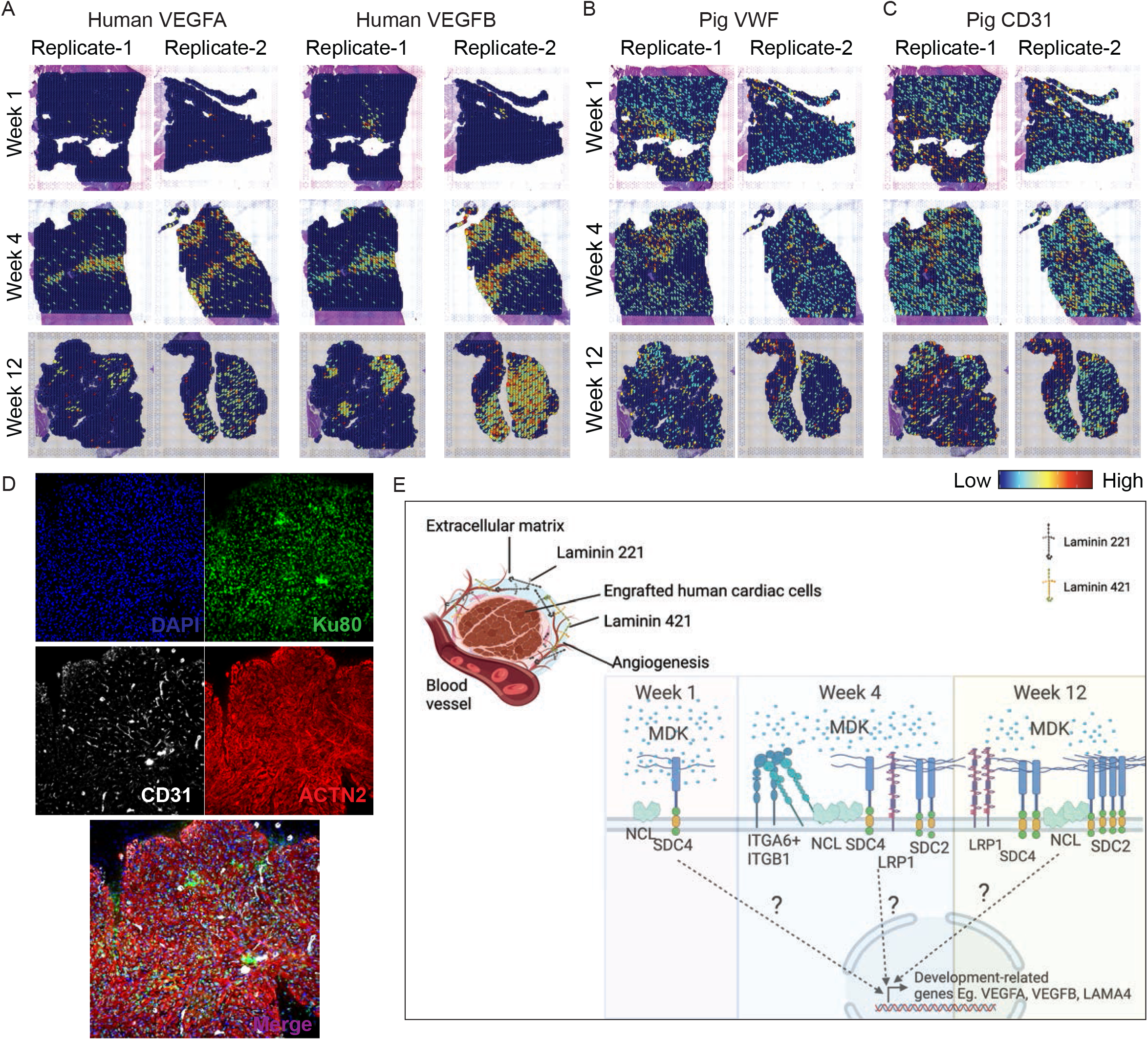
Expression of Vascular-related genes. **A-D)** H&E and spot overlays depict the normalized gene expression for human *VEGFA, VEGFB,* and pig *VWF and CD31* at 1-, 4- and 12- weeks post-transplantation. **(D)** Immunofluorescence staining of 12-weeks post-transplantation grafts with *CD31* and *ACTN2* antibodies. Human cells were identified using a human-specific Ku80 (huKu80) antibody. All nuclei are stained with DAPI (blue). **(E)** Illustration of the possible spatial orientation of laminin-221, laminin-421, and MDK signalling pathways towards angiogenesis in the human cells engrafted niche.

To confirm the formation of blood vessels, an immunohistology staining of the human graft at 12 weeks post-transplantation in the CMI model shows the expression of cardiac actinin-alpha-2 (ACTN2) and blood vessel-related CD31 protein (**Figure 6D**). This validated our ST findings of blood vessel regeneration in the engrafted human graft. We then speculated the mechanism of angiogenesis in this study which will enable future treatment to enhance the survival and regeneration capacity of engrafted CVP for ischemic heart patients. Figure 6E showed an illustration to summarise our findings where the engrafted human cells start maturing into functional CMs the pig’s infarcted heart. During the regeneration process, the expression of human MDK binds to its cell-cell surface receptors which resulted in a downstream signaling mechanism that could increase the expression of vascular-related genes such as *VEGFA* and *VEGFB* to attract the host endothelial cells to form collateral blood capillaries. This is an important process to ensure that the human graft survives long-term in an ischemic region.

In conclusion, we observed expression of the human MDK-receptor signaling pathway, both human and pig vascular-related genes (human: *VEGFA* and *VEGFB*, pigs: *VWF* and *CD31*) which could potentially contribute to blood vessels formation in the engrafted human spots.

## DISCUSSION

In the present study, we showed the importance of understanding the spatial context of gene expression and molecular mechanisms in myocardial infarction^46–48^. The studies utilized spatial transcriptomes to explore the transcriptional signatures and diversity across the infarction zone, the border zone, and the remote zone in the human post-MI hearts. This demonstrated the feasibility of using ST in gaining further insights into stem cell therapies in the regeneration of infarcted hearts. However, not many studies have utilized the ST pipeline in attempts to understand the mechanisms of engraftment, maturation, and regeneration of transplanted cells. Following our previous studies, where we established functional heart improvement and reduction in ventricular arrhythmia from the transplantation of human CVP^10, 11^, we performed a time-series ST on both AMI and CMI pig hearts. These analyses characterized the transplanted human cells in a spatial-temporal fashion to gain insights into the possible regeneration mechanism.

We captured ST of timepoints AMI 1- and 2- weeks and CMI 1-, 4- and 12- weeks post-transplantation. Briefly, our computational analyses revealed successful transplantation of human CVPs into MI pig hearts in both MI models. Our data indicated high reproducibility of transplanted human cells across time points. ST also enabled us to characterize the neighboring spots that are adjacent to the transplanted human cells in the AMI model. From the CMI model, we observed higher expression of mature cardiac markers (such as *ACTN2, CD36, ESRRA, MYH7, MYL2, TNNT2*, and *TNNI*). We also detected increased mitochondrial, ribosomal activities, calcium and potassium channel genes, and connexin-43 expression over time suggesting maturation of the human graft.

Researchers in the stem cell field agree that in vitro cultured hPSC-CMs are immature and there is active research in developing ways to mature these fetal-like CMs into adult-like CMs. Methods such as prolonged cell culture time, the addition of biochemical cues, modification to extracellular matrices, and 3-dimensional culturing^49^. Instead of maturing cells *in vitro*, *in vivo,* environment has been shown to be possible by providing all the necessary signals to mature neonatal CMs in the rat’s heart^50^. Similarly, to others, we have shown the expression of mature cardiac genes (*MYH7, MYL2, TNNI3*), ion channels (*CACNA1C, RYR2, KCNH2*), gap junctions (*GJA1*), mitochondria, and metabolic (*ESRRA*) genes in the pig hearts. Interestingly in our data, there was a sarcomeric isoform switching between human *MYH6* and *MYH7* further confirming the maturation of human cells^16^. Indicating the continual maturation of the transplanted human cells *in vivo*. However, whether these cells are more mature than what others have reported, detailed isolation and characterization of the live human cells after transplantation will be required.

In addition, we explored conserved human spot communication at the regions of transplantation which revealed MDK signaling. Previous studies have shown that the over-expression of MDK led to severe cardiac hypertrophy and adverse remodeling^51^. However, a moderate expression of MDK is shown to be necessary for arteriogenesis in collaterals^45^. Coincidentally, our data also revealed higher expression of MDK at 1- week which then gradually decreased and was maintained at 12- weeks suggesting the stimulation of vasculature without the adverse effects associated with the elevated level of MDK. The observed expression of vascular-related genes (*VEGFA, VEGFB, VWF, CD31*), as well as endothelial-related laminin alpha 4 (LAMA4)^52, 53^ (**Supplementary Figure 6**), suggested arteriogenesis for collateral blood vessels formation in the human graft.

Since laminins are important molecules in the extracellular matrix and the CVPs are differentiated on laminins, we explored the various laminins’ expressions. Considering all the laminin isoforms (*LAMA1, LAMA2, LAMA3, LAMA4, LAMA5, LAMB1, LAMB2, LAMB3, LAMB4 LAMC1, LAMC2* and *LAMC3*), we could only detect significant changes in *LAMA2, LAMA4, LAMA5, LAMB1, LAMB2,* and *LAMC1* genes (Supplementary Figure 6). Since laminins are structurally heterotrimeric containing alpha, beta, and gamma chains, we hypothesized that LN221, LN421, and LN521 are the most abundantly expressed isoforms in the heart muscle. It was known that cardiac-promoting laminin alpha 2 (*LAMA2*) is related to cardiac development^11^, laminin alpha 4 (*LAMA4*) is important in vasculogenesis and endothelial differentiation^52^, and laminin alpha 5 (*LAMA5*) is involved in embryogenesis and stem cells maintenance^54^.

It is interesting to compare our angiogenic findings with those of others who have transplanted cardiac-related cells into infarcted animal models. Chong *et al* have shown in non-human primates that by using microcomputed tomography, observed the host vessels’ perfusion into the transplanted hESC-CM graft^55^. Other reports by Romagnuolo *et al* showed the presence of high-density porcine microvessels within the hESC-CM graft using immunostaining^56^ and Laflamme *et al* also showed in rat hearts the few human-derived endothelial cells in contrast to host-derived endothelial cells^57^. Interestingly, Fernandes *et al* showed in hESC-CVP transplanted rat MI model the rare tissue staining of human-derived CD31 expression although the study and did not measure an increase in overall cardiac vascularization^58^. Taken together and placed into the context of transplanting human-derived cells into an infarcted preclinical animal model, we propose a vascularization mechanism of MDK-mediated VEGF host blood vessel formation from the spatiotemporal transcriptomic analysis. This process promoted graft survival and maturation leading to functional cardiac improvement and regeneration.

There are, however, limitations in our study and longer time point follow-up beyond 12- weeks would be beneficial to follow the long-term maturation of a human cardiac graft. We also know that the 10x ST technology is not at single-cell resolution, so our cell-to-cell communication could be improved as the resolution of spatial transcriptomics evolves. In our future study, we will be considering multiplexing spatial proteomics to allow high-resolution staining of multiple proteins such as cardiac-related genes, human Ku80, VEGFA, VEGFB, and MDK-receptor, which could be stained and imaged together for high spatial phenotyping. Besides that, we are also considering isolating the engrafted human cells from the infarcted pig tissues at various time points for electrophysiology studies, and further characterization to compare the maturity of these *in vivo* matured CMs as compared to *in vitro* matured CMs in the future study.

In summary, this paper describes the utilization of ST to study the spatiotemporal transcriptomic landscape of hPSC-derived CVPs transplanted into AMI and CMI pig hearts. We can differentiate both pig and human gene expression from each spot and identify with confidence the spots containing human cells. Further analysis with cell-to-cell communication identified ligand-receptor pairs (MDK signaling) that are upregulated across the time points to potentially induced angiogenic-related genes for the development of a pig’s collateral blood vessels. This mechanism is important for the survival and maturation of human cells *in vivo* with the impact on successful CVP-based regenerative cardiology.

## Acknowledgments

We would like to acknowledge Jing Guo for the constructive discussion on the ST analysis data. This work has been supported by grants from the NMRC and NRF of Singapore (MOH-STaR18may-0001 and CRP24-2020-0083) and Goh Cardiovascular Research (Duke-NUS-GCR/2020/0018).

## Author Contributions

This study was conceptualized and study designs were planned by S.A. and L.Y. The animal experiments, tissue processing, and histology were performed by S.L., Y.L., V.R., Y.W., R.T., C.T., and L.Y.C. The ST computational analyses, data interpretation, and visualization were performed by S.A. E.P. provided bioinformatics input and discussion to the manuscript. L.Y. and S.A. wrote the manuscript which was reviewed and edited by E.P. and K.T. L.Y. supervised the project, and funding was acquired by K.T. and L.Y.

## Supplementary Figures

**Supplementary Figure 1.**
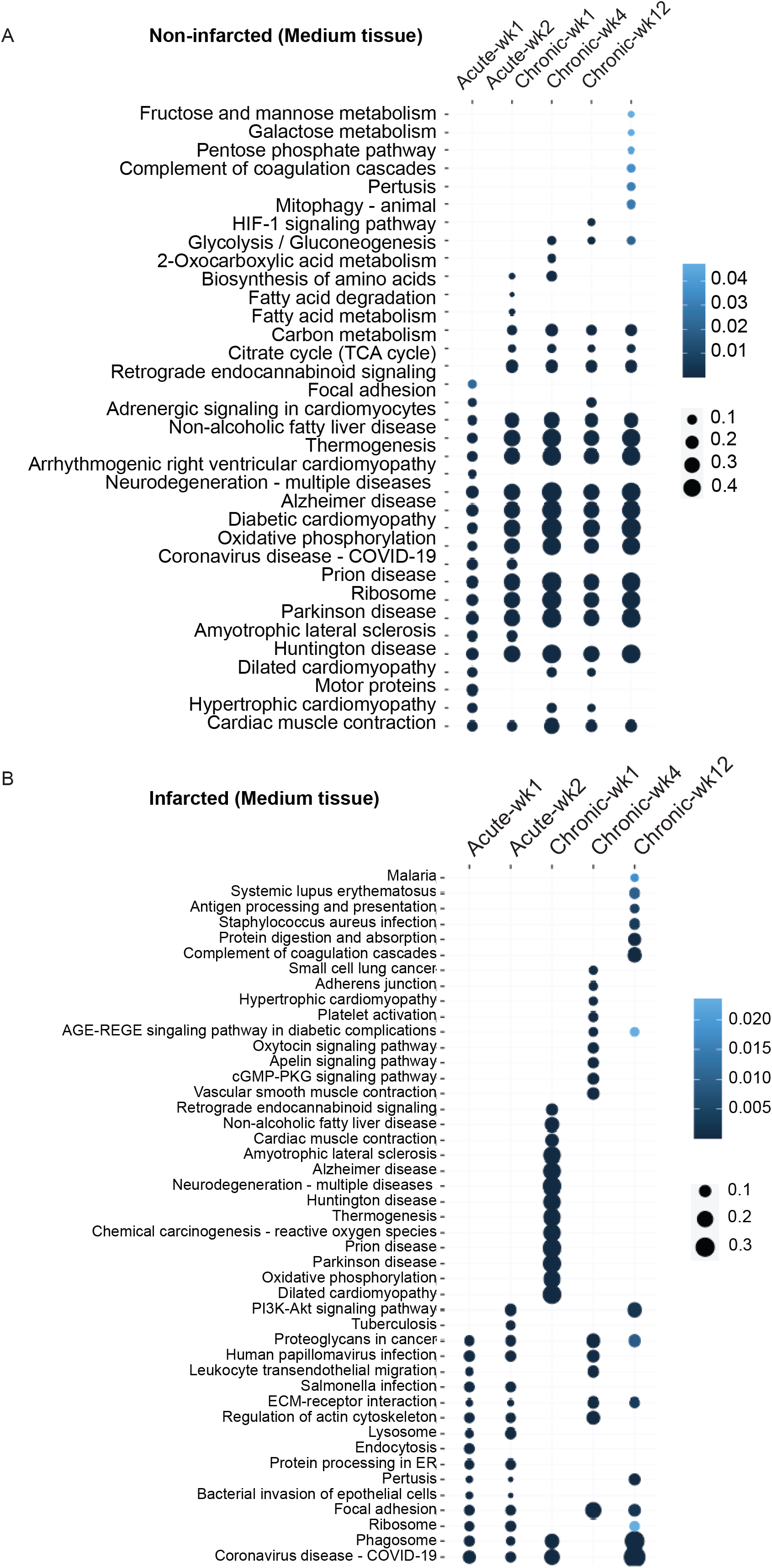
Functional enrichment analyses of NI and infarcted regions in medium tissues. **A-B)** Dotplot of pathways enriched for genes upregulated in **(A)** NI and **(B)** infarcted clusters. The dots are colored by p-values (FDR). The x-axes indicated the MI model and time points. GeneRatio is the percentage of the genes enriched in each pathway from NI and infarcted region against the total number of genes in the network.

**Supplementary Figure 2.**
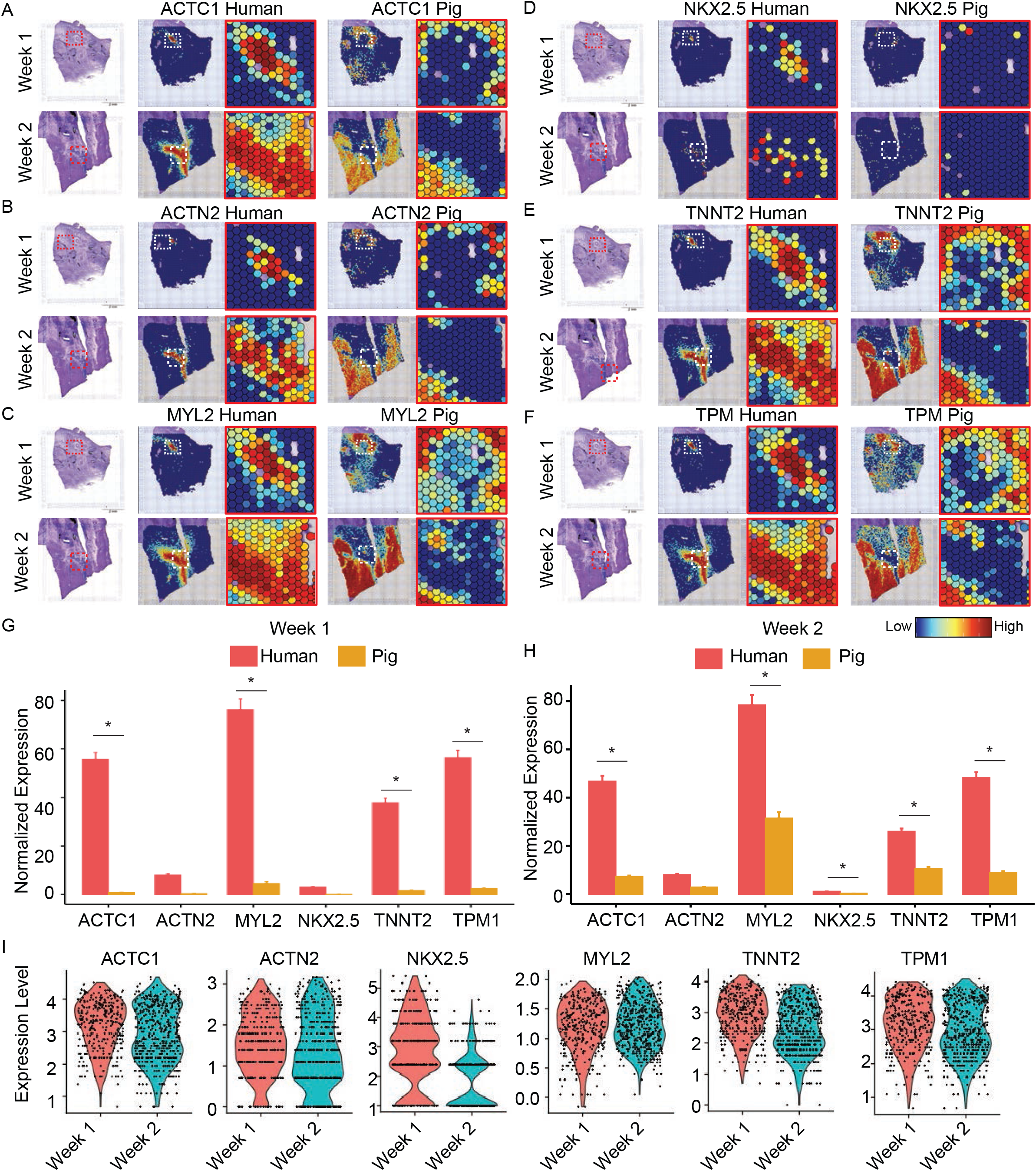
Characterization of Engrafted Human Cells (Replicate-2) in AMI Pig Hearts. **A-F)** H&E and spot overlays depict the normalized expression for human and pig *ACTC1, ACTN2, MYL2, NKX2-5*, *TNNT2,* and *TPM1* marker genes at 1- and 2-weeks post-transplantation. (Left) H&E images with engrafted human cells are highlighted in red. Insert shows the higher magnification of engrafted human cells from the dotted white box. **(G-H)** Quantification of the expressions of pig and human genes in the spots engrafted with human cells as shown in the red box highlighted in panels A-F in 1- and 2- weeks (* p-value < 0.05). **(I)** Violin plots showing the normalized expression of *ACTC1, ACTN2, MYL2, NKX2-5*, *TNNT2,* and *TPM1* marker genes across all time points.

**Supplementary Figure 3.**
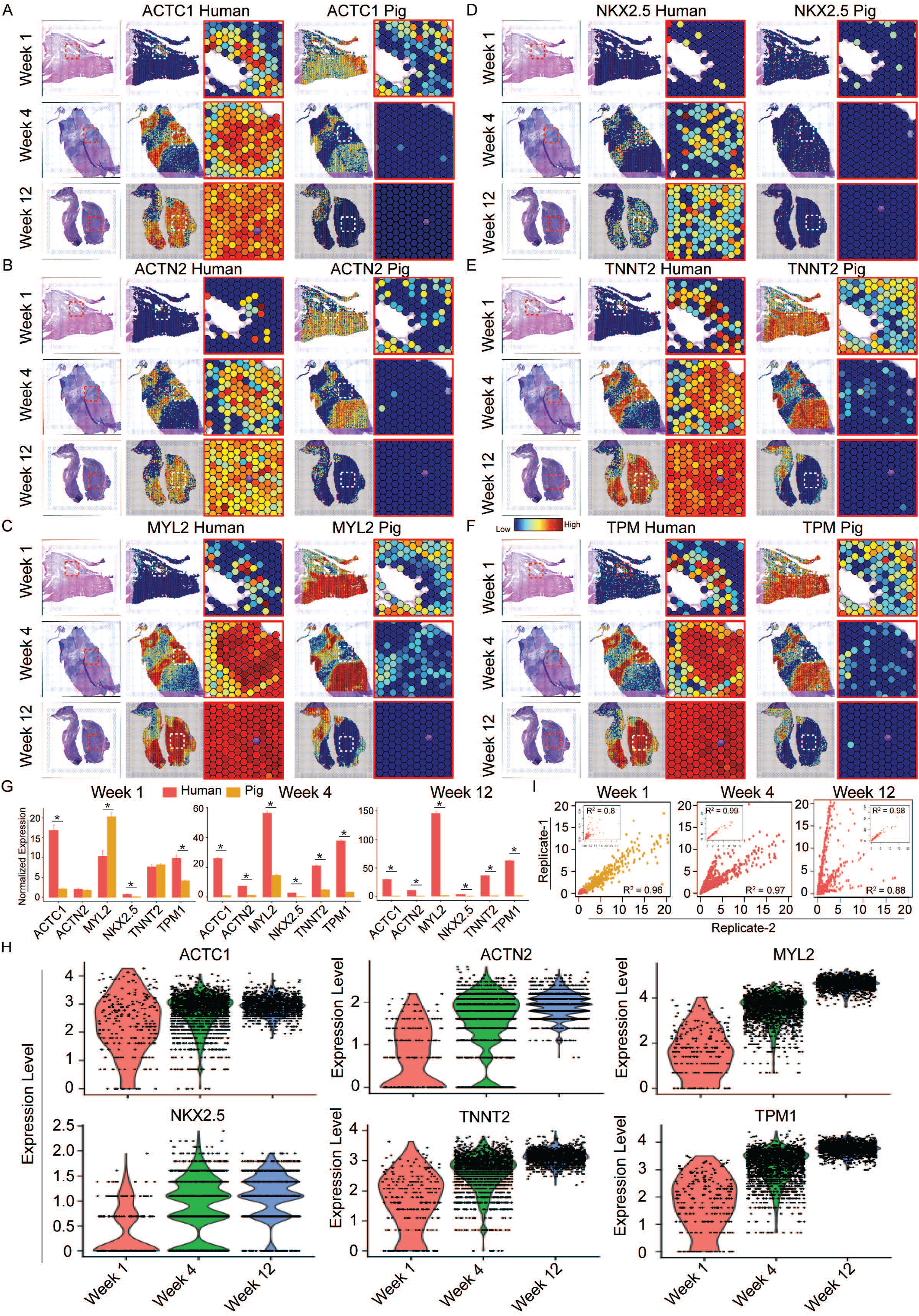
Characterization of Engrafted Human Cells (Replicate-2) in CMI Pig Hearts. **A-F)** H&E and spot overlays depict the normalized expression for human and pig *ACTC1, ACTN2, MYL2, NKX2-5*, *TNNT2,* and marker genes at 1-, 4- and 12-weeks post-transplantation. (Left) H&E images with engrafted human cells are highlighted in red. Insert shows the higher magnification of engrafted human cells from the dotted white box. **(G)** Barplots show quantified expressions of pig and human genes in the spots engrafted with human cells as shown in the highlighted panels A-F each week (* p-value < 0.05). **H)** Violin plots showing the normalized expression of *ACTC1, ACTN2, MYL2, NKX2-5*, *TNNT2,* and marker genes across all time points. **(I)** Correlation plot of all genes including pig and human genes from both replicates engrafted with human cells (R^2^ = 0.96, 0.97, and 0.88 at 1-, 4- and 12-weeks respectively). Insert shows the correlation of human genes only in engrafted regions of both replicates (R^2^ = 0.8, 0.99, and 0.98 at 1-, 4- and 12- weeks).

**Supplementary Figure 4.**
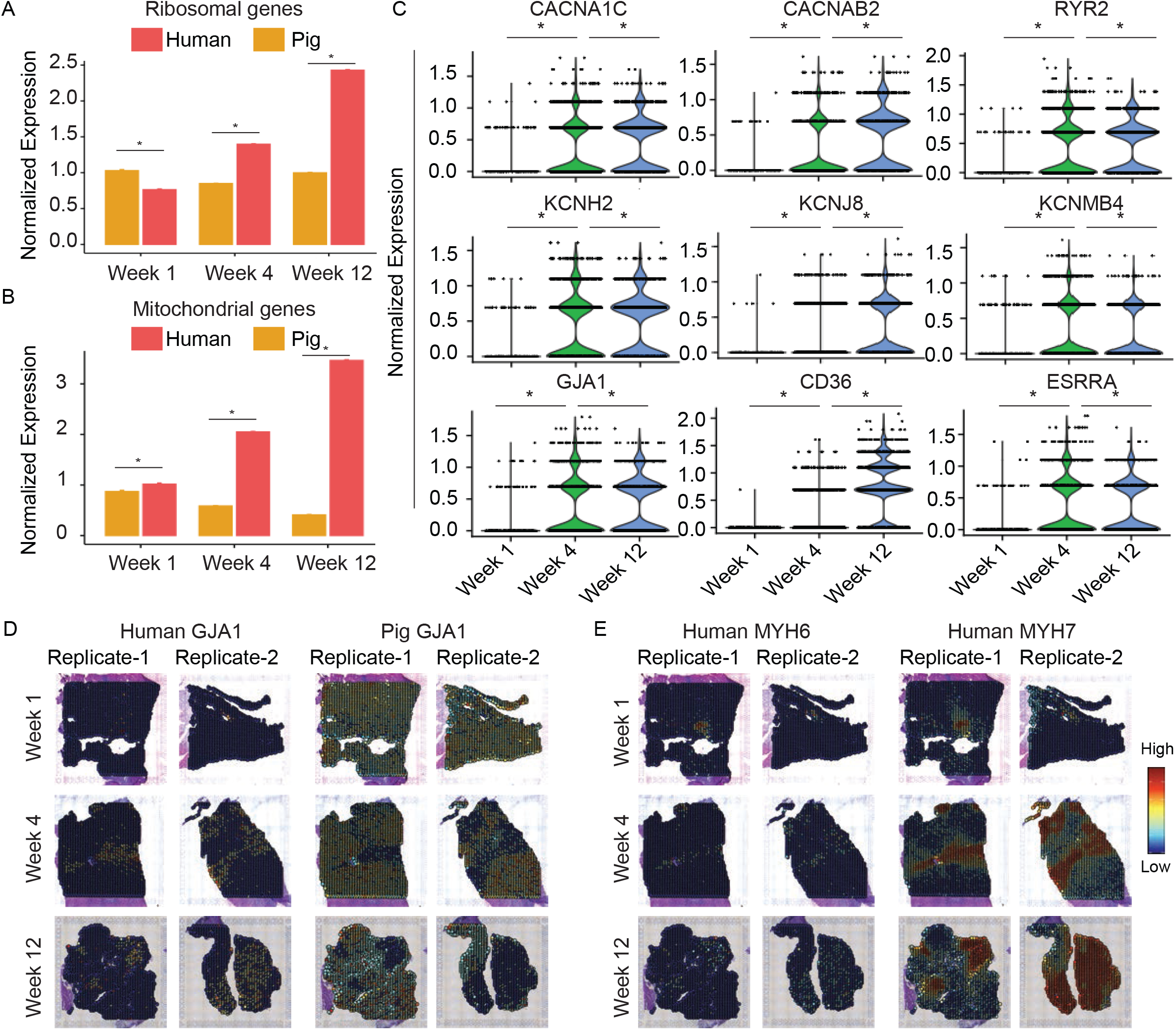
Metabolic Maturation of Human Grafts. **A-B)** Barplots show quantified expressions of pig and human **(A)** ribosomal and **(B)** mitochondrial genes in the spots engrafted with human cells as shown in the highlighted H&E staining in Figure 2 A-B (* p-value < 0.05). **C)** Violin plots showing the normalized expression of cardiac maturations genes *CACNA1C, CACNB3, RYR2, KCNH2, KCNJ8, KCNMB4, GJA1, CD36,* and *ESRRA* across all time points (* p-value < 0.05). **(D)** H&E and spot overlays depict the normalized expression for human and pig *GJA1* across all time points. **(E)** H&E and spot overlays depict the normalized expression for human *MYH6* and *MYH7* across all time points.

**Supplementary Figure 5.**
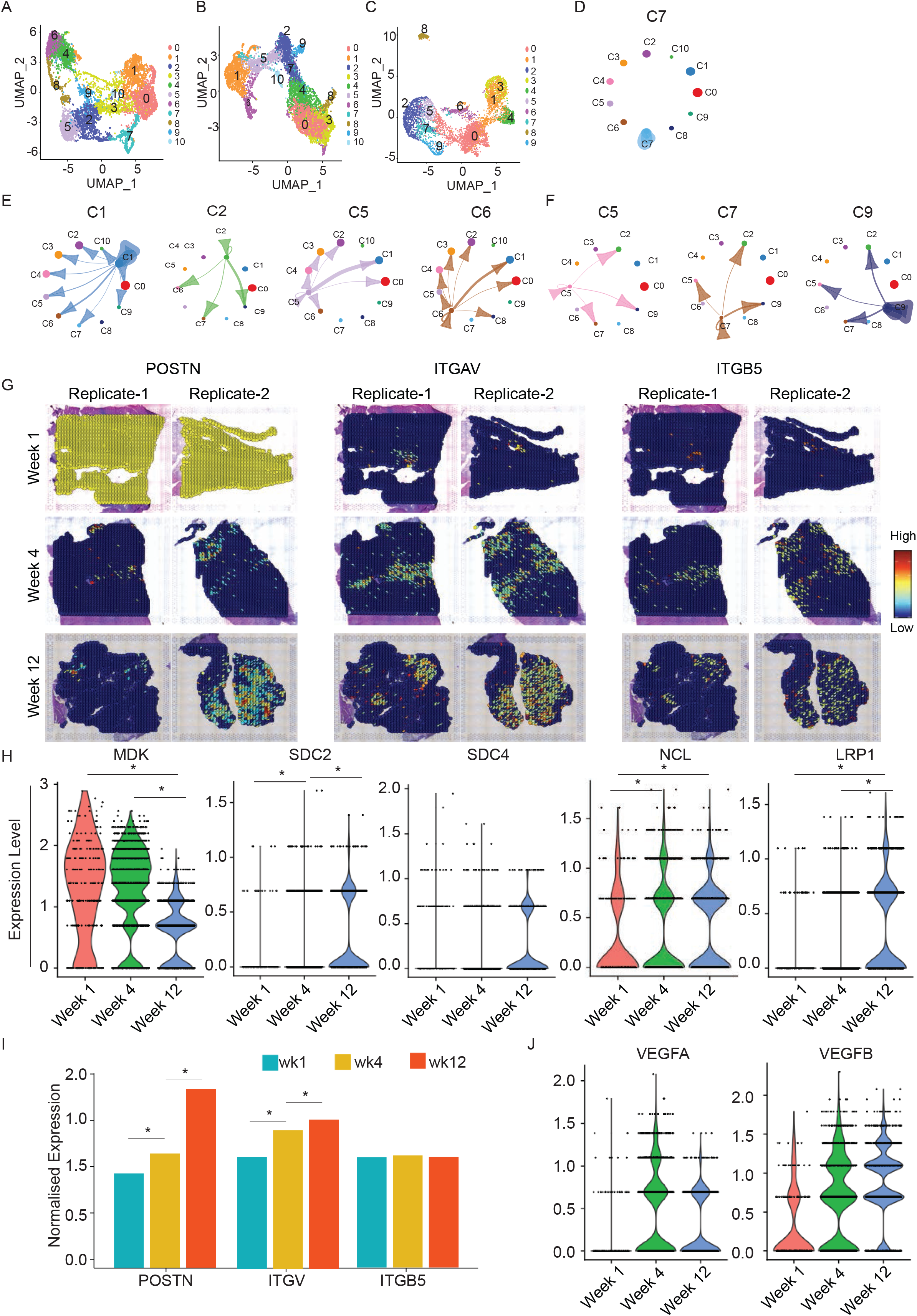
Cell-cell communications in human CVPs engrafted clusters in the CMI model. UMAPs showing the unsupervised clustering of spots based on global gene expression within individual spots for biological replicates of **(A)** 1- week **(B)** 4- weeks and **(C)** 12- weeks post-transplantation. Circle plots showing interactions with the ligand receptors within and across human clusters at **(D)** 1-, **(E)** 4- and **(F)** 12- weeks. The arrows indicate incoming or outgoing signalling at each cluster. The round loops at clusters represent the interactions within the same cluster. **(G)** H&E and spot overlays depict the normalized expression for human *POSTN*, *ITGAV,* and *ITGB5* at 1-, 4- and 12- weeks post-transplantation. **(H)** Violin plots showing the normalized expression of *MDK, SDC2, SDC4, NCL,* and *LRP1* marker genes across all time points (* p-value < 0.05). **(I)** Quantification of the expressions of *POSTN*, *ITGAV,* and *ITGB5* genes in spots engrafted with human cells across the time points (* p-value < 0.05). (J) Violin plots showing the normalized expression of *VEGFA* and *VEGFB* marker genes across all time points (* p- value < 0.05).

**Supplementary Figure 6.**
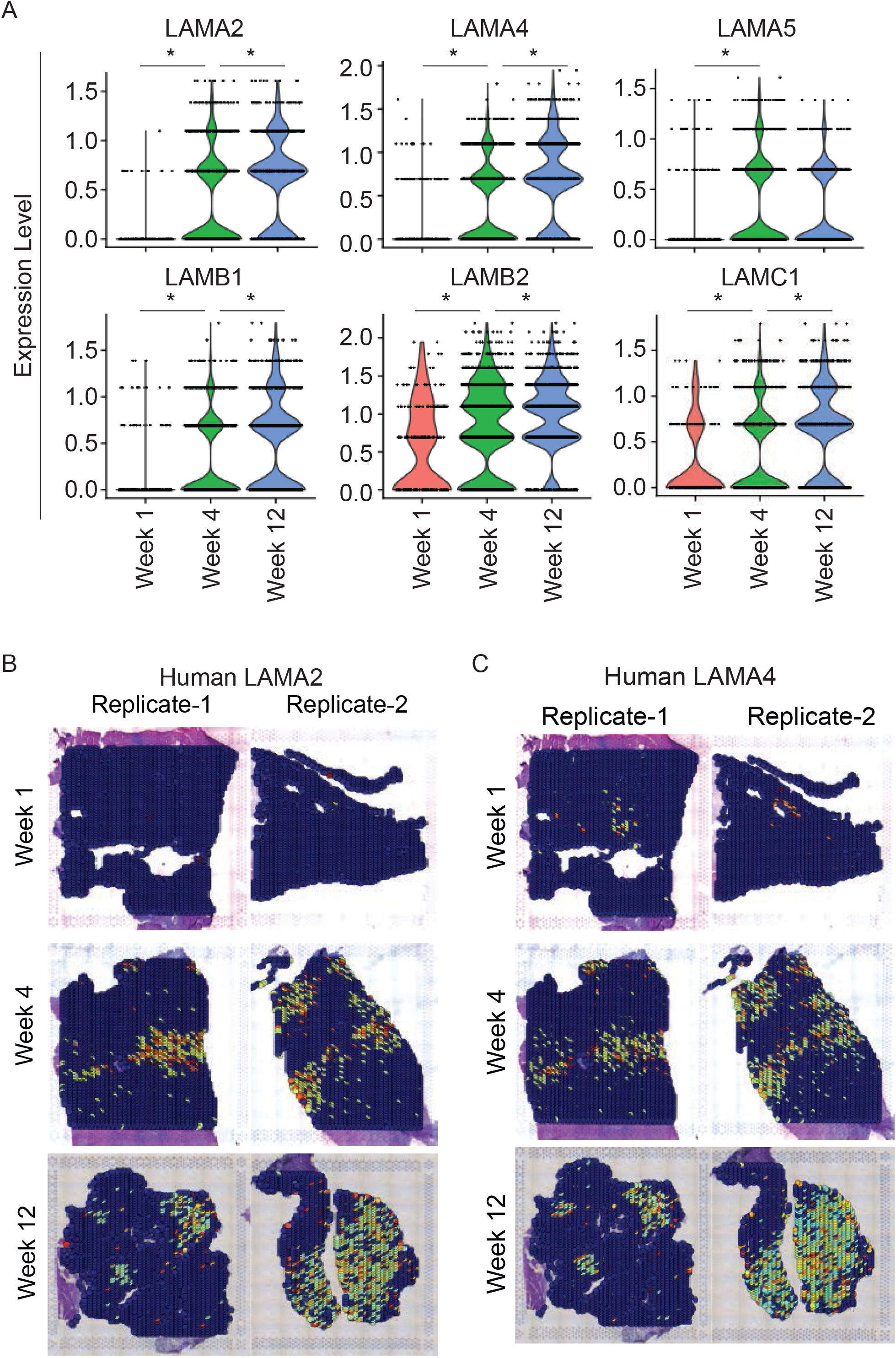
Expression of human laminin isoforms in CMI model. A) Violin plots showing the normalized expression of *LAMA2, LAMA4, LAMA5, LAMB1, LAMB2* and *LAMC1* marker genes at 1-, 4- and 12-weeks post-transplantation. (* p-value < 0.05). **(B-C)** H&E and spot overlays depict the normalized expression for human cardiac laminin isoforms *LAMA2* and *LAMA4* across all time points.

